# A patient-derived xenograft model of *FUS::TFCP2* intraosseous rhabdomyosarcoma reveals chemoresistance and differential sensitivity to ALK inhibitors

**DOI:** 10.64898/2026.08.05.743043

**Authors:** Shefali Chauhan, Kristen Jones, Victoria Krajbich, Barron Smith, Cole McCallister, Tiffany Bui, Rebecca Smith, Randy L. Woltjer, Sintawat Wangsiricharoen, Duncan C Ramsey, Monika A. Davare

## Abstract

*TFCP2*-rearranged rhabdomyosarcoma is an exceptionally rare and highly aggressive malignancy driven by TFCP2 gene fusions and associated with a dismal clinical prognosis. Because standardized treatment regimens are lacking, developing representative preclinical models is critical for identifying effective therapies. Here, we present a case of a 29-year-old male with rapidly progressive, metastatic pelvic intraosseous rhabdomyosarcoma (iRMS) harboring a FUS::TFCP2 fusion and anaplastic lymphoma kinase (ALK) overexpression. To evaluate therapeutic vulnerabilities, we established a patient-derived xenograft (PDX) model that faithfully recapitulated the histologic, immunohistochemical, and molecular hallmarks of the primary tumor. High-throughput in vitro pharmacological screening of PDX-derived cells demonstrated notable resistance to standard cytotoxic chemotherapies and revealed a paradoxical and selective sensitivity profile across ALK inhibitors. The PDX-derived cells were susceptible to crizotinib, brigatinib, and ceritinib, yet resistant to the more selective second- and third-generation inhibitors alectinib and lorlatinib. Notably, next-generation ROS1/pan-TRK inhibitors (entrectinib, repotrectinib, and taletrectinib) demonstrated superior efficacy compared to the fourth-generation ALK inhibitor NVL-655. Our findings establish a validated preclinical PDX model for FUS::TFCP2 iRMS and suggest that multi-targeted tyrosine kinase inhibition may offer a more viable therapeutic strategy than narrow-spectrum ALK targeting or conventional chemotherapy.

## Introduction

Rhabdomyosarcoma (RMS) is a malignant soft-tissue tumor arising from primitive mesenchymal cells of striated muscle lineage ^1 2 3^. Based on histopathology, RMS has been classified into four subtypes: alveolar, embryonal, spindle/sclerosing, and pleomorphic ^4^. The 2020 World Health Organization (WHO) classification recognized that a subset of spindle cell/sclerosing RMS can affect bone as in this case of iRMS^5^.

Intraosseous RMS follows a highly aggressive clinical course ^4^. The majority (~70%) arise in the craniofacial bones, particularly the maxilla and mandible ^6^, with the remainder usually occurring in the pelvis, femur, or vertebrabone. iRMSs harbor recurring gene fusions, most commonly *EWSR1/FUS::TFCP2* and less commonly, *MEIS1::NCOA2* ^7 8 9 10 11^.

Histologically, *EWSR1*/*FUS*::*TFCP2* iRMS is composed of spindle and epithelioid cells arranged in sheets or fascicles with abundant eosinophilic cytoplasm. The nuclei are irregular and hyperchromatic with prominent nucleoli. The stroma is variable and often contains focally sclerotic areas^12 7^. Immunohistochemistry reveals expression of skeletal muscle markers (desmin, MYOD1, myogenin), epithelial markers (pan-keratins), and ALK ^6 7^.

Previous genomic and transcriptomic analyses have demonstrated high expression of ALK and Telomerase Reverse Transcriptase (TERT) and homozygous deletions of *CDKN2A* and *MTAP* (seen in ~75% of cases) in a background of high genomic instability ^13 14^. *TFCP2*-fusion RMSs remain poorly understood, and no standard treatment regimen has been established. While platinum-based chemotherapeutics and *ALK* inhibitors have shown some promise in limited studies, the overall prognosis remains dismal, with a reported median overall survival of approximately 29 months ^13-15^.

This report presents a case study of a 29-year-old male with pelvic *FUS*::*TFCP2* iRMS. Because standardized treatment regimens are lacking, developing representative preclinical models is critical for identifying effective therapies. To investigate the therapeutic vulnerabilities of this rare malignancy, we established a patient-derived xenograft (PDX) model and utilized PDX-derived cells to conduct functional *in vitro* drug sensitivity testing, comparing the efficacy of cytotoxic therapies against an array of tyrosine kinase inhibitors with activity against ALK.

## Materials and Methods

This study was approved by the Oregon Health & Science University Institutional Review Board (IRB#26657). Clinical information was obtained from the medical records and from the pathologists and clinicians.

### Molecular Profiling and Next-Generation Sequencing

Molecular profiling was performed using the GeneTrails® Solid Tumor Panel at the Knight Diagnostic Laboratories (Oregon Health & Science University), as previously described ^16,17^. Briefly, targeted amplicon-based libraries generated from tissue-extracted genomic DNA were sequenced on the Ion Torrent Personal Genome Machine (Thermo Fisher Scientific, Waltham, Massachusetts) to detect single nucleotide variants and indels, with variant calling and clinical annotation conducted via Torrent Suite Software (v4.0). Concurrent targeted microarray analysis was performed using the CytoSure™ Consortium Cancer SNP array (Oxford Gene Technology. Kidlington, Oxford). Following hybridization and scanning (Agilent SureScan Dx, Santa Clara, California), data were analyzed using CytoSure Interpret Software (Human Genome Build [hg19]) to call copy number variants and regions of copy-number neutral loss of heterozygosity. The targeted RNA panel is designed to detect fusions involving the 43 genes (including *FUS*) and is agnostic with respect to fusion partners. The gene fusions can be detected to the range of approximately 1-5% of input cells.

### Immunohistochemistry

Clinical Sample: Immunohistochemical analysis of patient biopsy specimens was performed at OHSU (Portland, OR) on formalin-fixed paraffin-embedded (FFPE) tissue sections. Automated staining was conducted using Ventana automated immunostainers (Ventana Medical Systems, Oro Valley, Arizona) according to validated clinical protocols. The primary antibody panel included: desmin (dilution 1:50–1:100), myogenin (1:25–1:50), cytokeratin cocktail (AE1/AE3/PCK26, 1:100), ALK (Clone ALK1, 1:100), smooth muscle actin (1:50), and S100 protein (1:50).

PDX Sample: For Patient-Derived Xenograft (PDX) samples, FFPE sections (4 - 5 µm) were mounted on positively charged slides and baked at 60°C for 1 hour. Automated deparaffinization and rehydration were performed using a Tissue-Tek® DRS™ Slide Stainer (Sakura Finetek USA, inc., Torrance, CA) through a graded series of xylene and ethanol washes, terminating in distilled water.Heat-induced epitope retrieval (HIER) was performed in 10 mM sodium citrate buffer (pH 6.0) using a water bath at 80–85°C for 45 minutes.

Following a 20-minute cool-down at room temperature, sections were rinsed in deionized water and washed in PBS. Endogenous peroxidase activity was quenched by incubating slides in 3% H_2_O_2_ in methanol for 10 minutes at room temperature. To minimize non-specific background, sections were blocked with 3% non-fat milk in TBS containing 0.1% Tween-20 and sodium azide for 1 hour at room temperature.

Sections were incubated overnight at 4°C with primary antibodies diluted in 5% BSA-TBS containing 0.1% Tween-20 with sodium azide. Dilutions were as follows: 1:500 for mouse anti-pan-cytokeratin (AE1/E3) (MNF116, Dako, Carpinteria, CA 93013, M0821), rabbit anti-desmin (F5V4, Cell Signaling Technology, Danvers, MA 01923, USA), mouse anti-myogenin at 1:2000 (F5D, Dako, Carpinteria, CA 93013, M3559), and rabbit anti-SATB2 at 1:500 (EP281, Cell Marque, Rocklin, CA 95677, 384R-16); and 1:400 for anti-ALK (D5F3, Cell Signaling Technology). Following three washes in PBST, sections were incubated with an HRP-conjugated goat anti-rabbit or HRP-conjugated goat anti-mouse secondary antibodies (Vector Laboratories Inc., Newark, CA 94560, BA-1000, BA-2000), (1:200) for 1 hour at room temperature.

Signal was visualized using 3,3’-Diaminobenzidine (DAB) substrate (Vectastain ABC kit, Peroxidase (Standard), Vector Laboratories Inc., Newark, CA 94560), with reaction times monitored via microscopy (typically 3–10 minutes). Sections were counterstained with Hematoxylin, dehydrated via an automated Sakura Finetek program (Sakura Finetek USA, inc.), cleared in xylene, and permanently mounted using ( Cytoseal 60, Epredia, Kalamazoo, MI 49008). Stained sections were visualized using a ZEISS Axioscan7 slide scanner at the OHSU Advanced Light Microscopy Core (RRID:SCR_009961). The images were captured at 20x magnification.

### PDX tumor model

Fresh tumor tissues were obtained from the patient following informed consent under an Institutional Review Board (IRB)-approved protocol. Samples were collected aseptically and transported in ice-cold HBSS supplemented with 1% penicillin streptomycin and amphotericin B (Fisher Scientific). For long-term storage, fragments were preserved in Bambanker (FUJIFILM Biosciences, Santa Ana, CA) and stored in liquid nitrogen. To establish the PDX model, cryopreserved fragments were thawed rapidly and washed in PBS. Tumor fragments (approximately 3 to 4 mm) were implanted subcutaneously into the flank of 6–8 week old female NSG mice (n=2, The Jackson Laboratory). Implantation was performed under isoflurane anesthesia. Post operative analgesia was administered following implantation. Mice were monitored weekly for clinical health and tumor progression. Tumor volume (V) was calculated twice weekly from digital calipers using the standard formula V = (length x width^2^) / 2. Animals were euthanized via CO_2_ inhalation followed by cervical dislocation once tumors reached a volume between 1500 - 2,000 mm^3^ or if humane endpoints were met. All in vivo experiments were approved by the Oregon Health and Science University Institutional Animal Care and Use Committee (IACUC; Protocol # TR02_IP00001484) and conducted in accordance with the American Veterinary Medical Association (AVMA) guidelines. All studies utilized PDX passages 1–2 to ensure the maintenance of the primary tumor’s molecular architecture.

### RNA isolation and RT-PCR

Total RNA was extracted from the PDX-derived primary culture cells using the commercial RNA isolation kit, RNeasy® Mini Kit (Qiagen, Hilden, Germany) following manufacturer’s instructions and stored at −80°C. The RNA quantity and quality were assessed using NanoDrop spectrophotometer (Thermo Fisher Scientific).

For RT-PCR, cDNA was generated from 100 ng of RNA, using SuperScript™ IV VILO™ Master mix with ezDNase (Invitrogen), following the manufacturer’s instructions. PCR amplification of cDNA was carried out in a 20 µL volume containing Platinum™ SuperFi II PCR Master Mix] (Invitrogen) and set of primers. The cycling conditions were initial denaturation at 98°C for 2 minutes, followed by 40 cycles of (denaturation at 98°C, 10 seconds), (annealing at 58.2°C, 10 seconds), (extension at 72°C for 30 seconds) with final extension at 72°C for 5 minutes. The PCR products were analyzed by electrophoresis on a 2% agarose gel stained with SYBR™ safe (Invitrogen).

### Western Blot

PDX-derived cell lines were harvested at 80% confluence and washed twice with ice-cold PBS. Total protein was extracted using cell lysis buffer (0.1% SDS, 0.5% sodium deoxycholate, 1% NP-40, 50 mM Tris-HCl pH 7.4, 150 mM NaCl, protease and phosphatase inhibitors). Protein concentrations were determined using the Pierce™ BCA Protein Assay (ThermoFisher Scientific). Samples containing 20 µg of total protein were denatured in Laemmli sample buffer supplemented with 2-mercaptoethanol for 10 min at 95°C and separated on 4–12% precast gradient Tris–glycine gels (Invitrogen; ThermoFisher Scientific). Proteins were transferred to nitrocellulose membranes (Prometheus, San Francisco, California). The membrane was blocked for 2 hours at room temperature in 5% Bovine Serum Albumin (BSA) in TBS-T (Tris-buffered saline with 0.1% Tween-20) to minimize non-specific binding, particularly for phosphate-specific detection. Membranes were incubated overnight at 4^0^C with primary antibodies, rabbit anti-ALK (clone D5F3, Cat#3633, Cell signaling Technology,1:2000), rabbit-anti-phospho-ALK (Cell Signaling, Cat# 3983S, Cell Signaling Technology, 1:1000), rabbit-anti-GAPDH (Cell Signaling, Cat#2118S, 1:10,000) diluted in 5% BSA/TBS-T. Following three 10-minute washes in TBS-T, membranes were incubated with HRP-conjugated secondary antibodies, anti-rabbit (Prometheus, 1:10,000) and anti-mouse (Prometheus, 1:10,000) for 2 hours at room temperature.

Chemiluminescent signal was generated using Clarity Western ECL Substrate (Bio-Rad, Hercules, California) and imaged using a ChemiDoc MP Imaging System (Image Lab / Image Studio™ Software v5.2)

### Dose Response Cell Viability Assays

Alectinib, ceritinib, crizotinib and NVL-655 were purchased from Selleckchem (Houston, TX). Methotrexate, gemcitabine and doxorubicin MedChem Express (Monmouth Junction, New Jersey). All other agents were purchased from Selleckchem.

All agents were received as a dried powder and were reconstituted in dimethyl sulfoxide (DMSO) (Sigma Aldrich, St Louis, MO) to a concentration of 10 mM or lower based on solubility specifications per manufacturer.

Cells were grown in T-75 flasks until 80% confluent, tryspinized, and plated at a density of 10,000 cells per well in a 384-well microtiter plate and incubated at 37°C with humidified 5% CO_2_ for 24 h. Agents were plated at 11 concentrations (10 µM, 4.38 µM, 1.92 µM, 0.84 µM, 0.37 µM, 0.161 µM, 0.071 µM, 0.031 µM, 0.014 µM, 0.006 µM and 0.003 µM) in triplicate using the HP Tecan D300e (Tecan, Männedorf, Switzerland). Cell viability was measured using Cell Counting Kit-8 (Selleckchem) per manufacturer’s instructions. Luminescence was measured using a BioTek Synergy H1 microplate reader (BioTek, Winooski, VT). IC_50_ values were determined using a nonlinear best fit in GraphPad Prism 10.6.0 (GraphPad, San Diego, CA).

## Results

### Clinical Presentation and Imaging

A 29-year-old male presented with three months of progressive and severe left hip and pelvic pain. Initial x-rays revealed a large destructive lesion of the left ilium (**Figure 1A**). Magnetic Resonance Imaging (MRI) demonstrated a heterogeneously enhancing mass with significant osseous destruction and soft-tissue extension (**Figures 1B, 1C**). Although initial staging computed tomography (CT) was negative for distant metastasis, a subsequent 18-Fluorodeoxyglucose Positron Emission Tomography (FDG-PET/CT) scan (**Figure 1D**) done one month later confirmed a large, highly metabolically active pelvic mass with intense peripheral tracer uptake (Standardized Uptake Value [SUV] max 15.7) with central necrosis and multiple FDG-avid bilateral pulmonary nodules consistent with metastasis.

**Figure 1.**
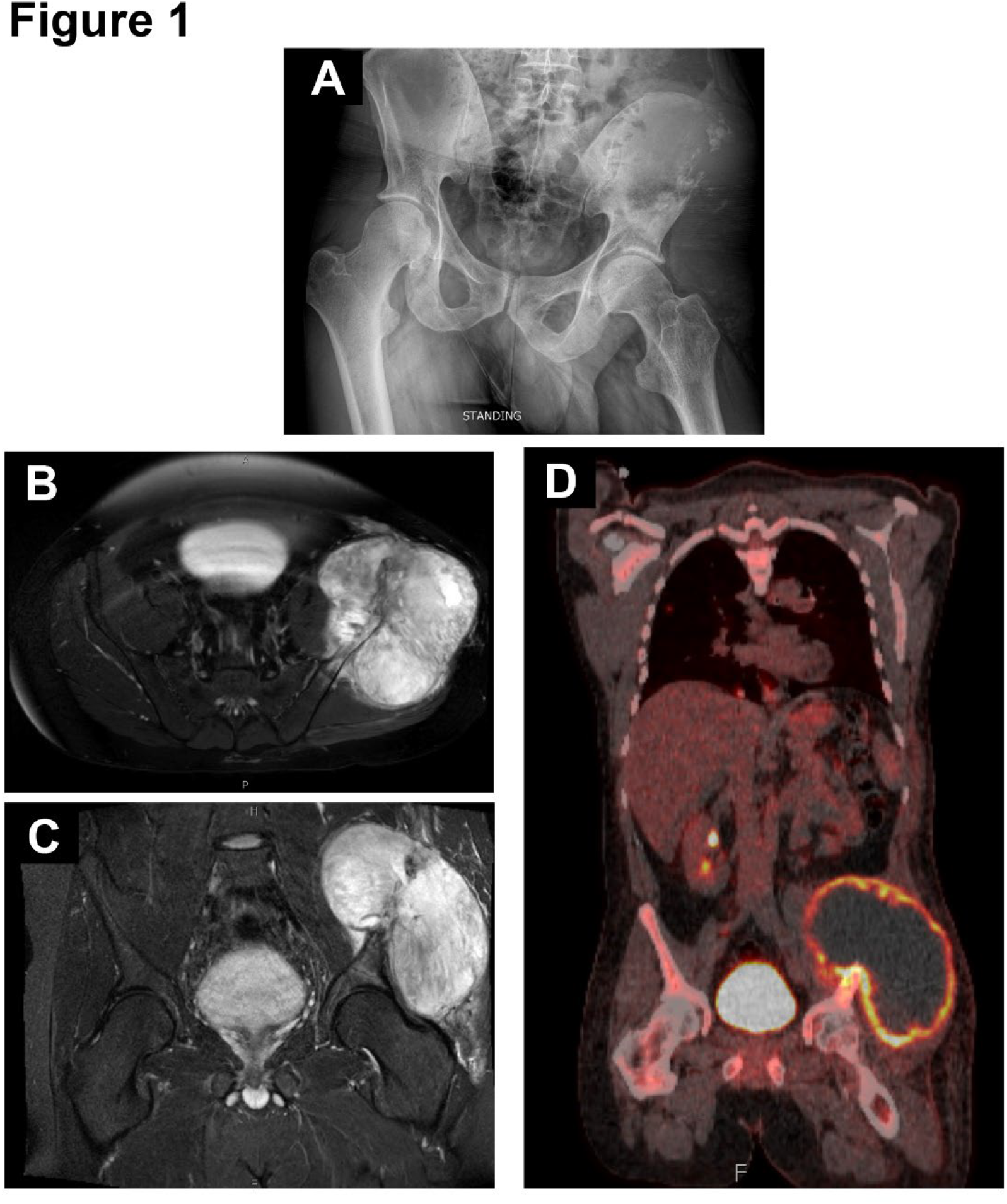
Clinical presentation and imaging of pelvic intraosseous rhabdomyosarcoma. (A) Radiograph of the pelvis showing a large destructive lesion of the left ilium. (B, C) Magnetic resonance imaging demonstrates a heterogeneously enhancing mass with extensive osseous destruction and soft-tissue extension. (D) 18F-fluorodeoxyglucose positron emission tomography/computed tomography (FDG-PET/CT) performed approximately one month later, showing a large, highly metabolically active pelvic mass with intense peripheral tracer uptake (SUVmax 15.7) and central necrosis, together with multiple FDG-avid bilateral pulmonary nodules consistent with metastatic disease.

### Immunohistochemical Profile and Lineage Characterization

Histologically, the tumor showed both epithelioid and spindle morphologies: one component consisted of sheets of epithelioid cells with irregular hyperchromatic nuclei and abundant eosinophilic cytoplasm, while other areas displayed spindled cells arranged in fascicules (**Figures 2A, 2B**). Extensive tumor necrosis and a high mitotic rate (11/10 high-power fields) were observed (**Figure 2C, 2D**).

**Figure 2.**
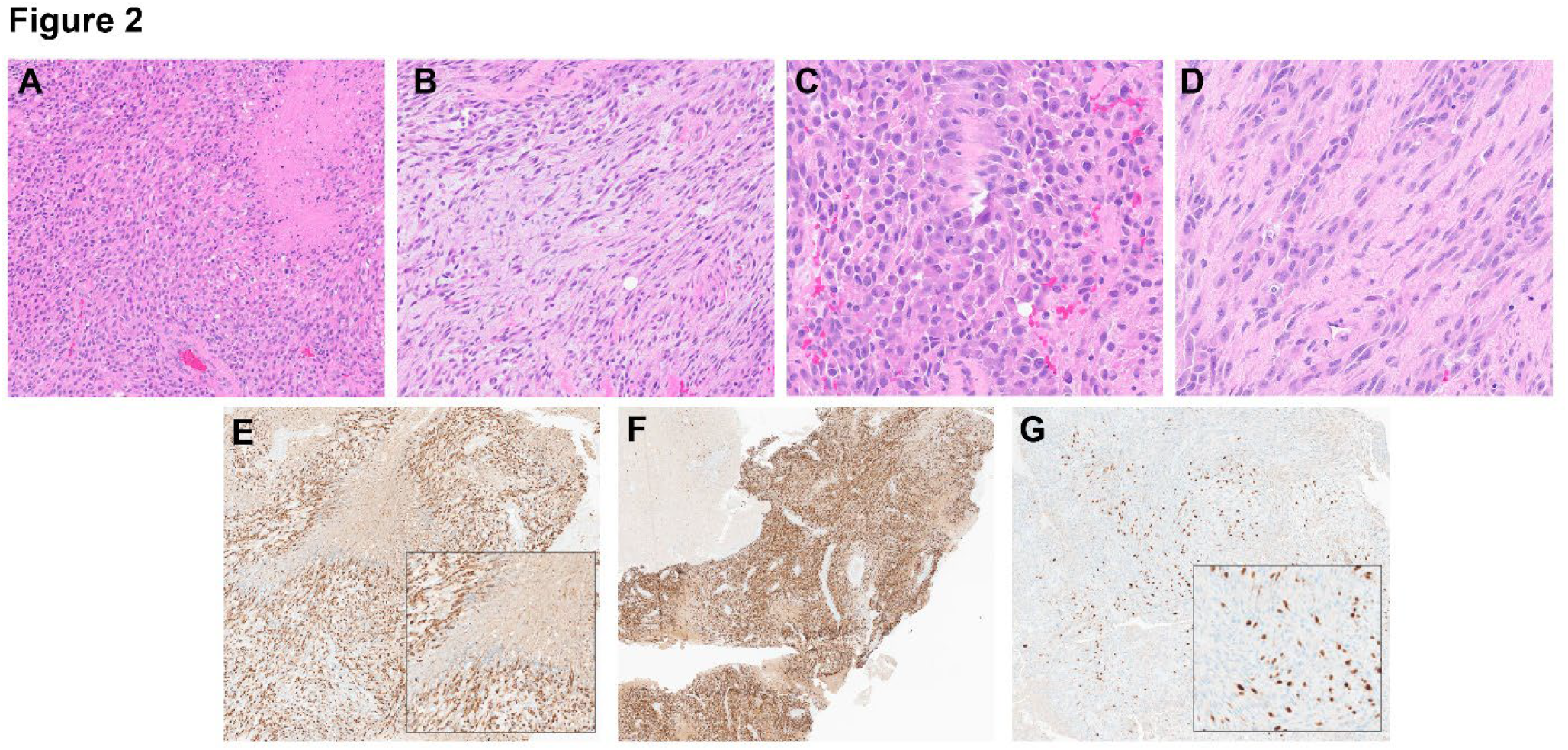
Histomorphology and immunohistochemical lineage characterization of the primary tumor. (A, B) Hematoxylin and eosin (H&E)-stained section showing two distinct morphologies: (A) sheets of epithelioid cells with hyperchromatic nuclei and abundant eosinophilic cytoplasm, and (B) spindled cells in a fascicular arrangement. (C, D) Extensive tumor necrosis (C) and frequent mitotic figures (D). (E) Strong, diffuse expression of pan-keratin AE1/AE3. (F) Strong, diffuse desmin expression. (G) Focal myogenin expression.

The tumor cells exhibited strong and diffuse keratin AE1/AE3 expression (**Figure 2E**) as well as labeling of rhabdomyogenic markers (diffuse desmin, diffuse myo D1, and focal myogenin expression) (**Figure 2F, 2G**). ALK was also diffusely positive.

### Molecular Profiling

RNA sequencing of the tumor sample, with estimated 80% tumor content at time of assay, revealed a chimeric mRNA transcript (chr16:31,196,500::chr12:51,512,555) representing an in-frame fusion between exons 1 to 6 of *FUS* and exons 2 to 15 of *TFCP2*. This rearrangement merges the N-terminal transcriptional activation domain of *FUS* with the CP2 DNA-binding and sterile alpha motif (SAM) domains of *TFCP2* (**Figure 3A**). The *FUS::TFCP2* fusion was independently confirmed by whole-genome sequencing.

**Figure 3.**
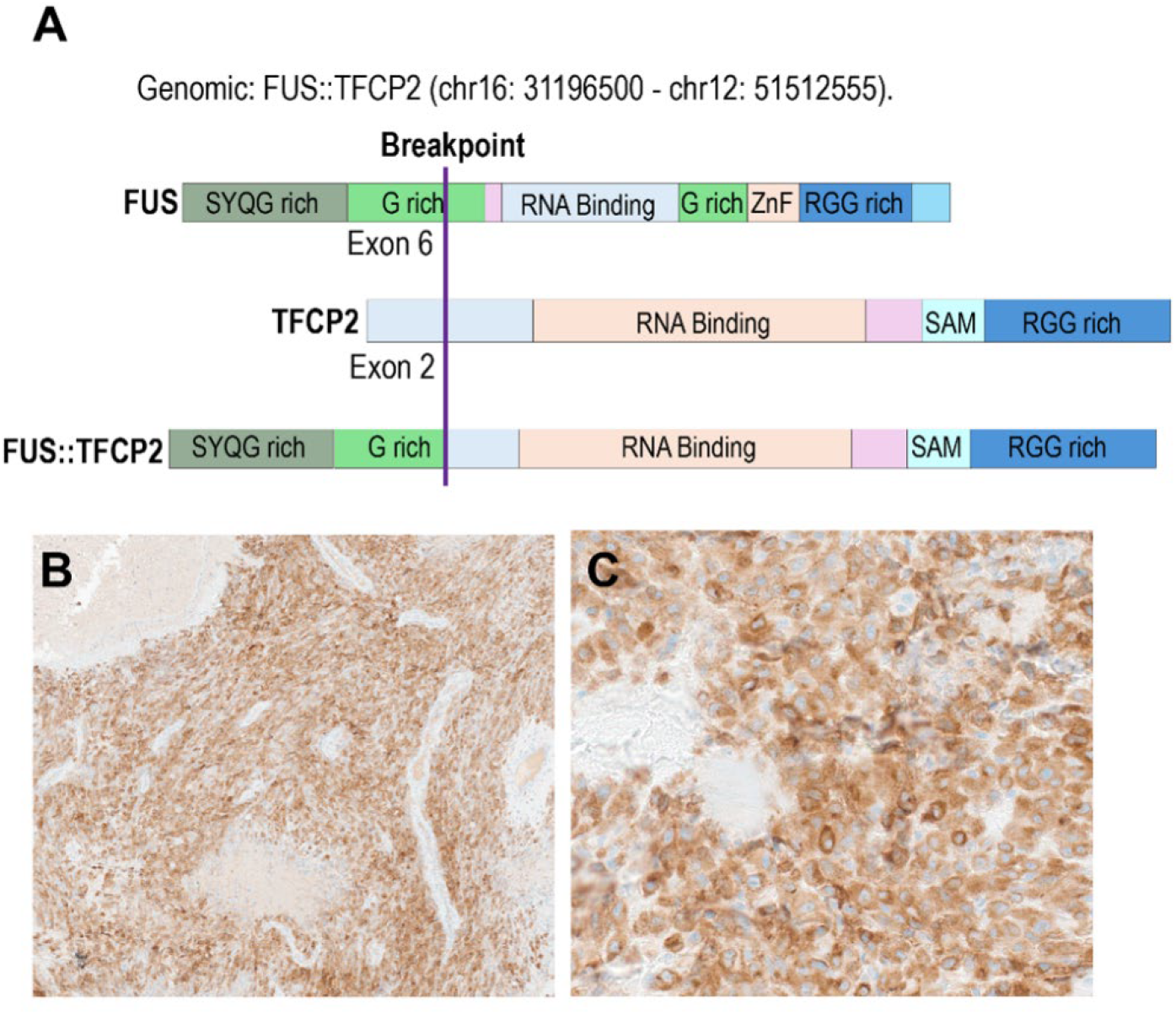
Identification of the *FUS::TFCP2* fusion and *ALK* overexpression. (A) Schematic of the in-frame FUS::TFCP2 fusion, joining exons 1–6 of FUS with exons 2–15 of *TFCP2*, fusing the N-terminal transcriptional activation domain of *FUS* to the CP2 DNA-binding and sterile alpha motif (SAM) domains of *TFCP2*. (B) Immunohistochemistry showing robust, diffuse cytoplasmic ALK expression in the tumor. (C) Further magnification of panel (B).

*FUS::TFCP2* fused oncogenes have been previously identified^11-15 18-20^. Consistent with the molecular profile of *FUS::TFCP2*-driven tumors, IHC evaluation demonstrated robust and diffuse cytoplasmic overexpression of ALK (**Figure 3B, 3C**). This ALK positivity serves as a diagnostic hallmark; when observed alongside skeletal muscle markers, it distinguishes *FUS::TFCP2*-driven iRMS from morphological mimics, including classical rhabdomyosarcoma and osteosarcoma.

Targeted DNA sequencing of the tumor (80% tumor content; mean coverage 1152x) identified no Tier I or copy-number alterations. The tumor was microsatellite stable (0 of 202 microsatellite sites abnormal; MSI-High threshold, >5%) with a low tumor mutational burden (2 mutations/Mb). Two variants of unknown significance were detected: *KIT* c.1430C>T (p.Ser477Phe) and *RB1* c.2263T>C (p.Phe755Leu). Neither represents a recognized activating or inactivating alteration, and both are reported as Tier III for completeness without established functional significance.

### Targeted Microarray Analysis Reveals Complex Chromosomal Imbalances and Copy-

#### Neutral Loss of Heterozygosity

Targeted oncology microarray was performed on the patient biopsy specimen to evaluate for broader genomic copy number variations. Although the interpretation of certain genomic regions was partially limited by probe scattering and single nucleotide polymorphism call inconsistencies, several significant copy number imbalances were confidently identified. These alterations included an interstitial deletion of on the long arm of chromosome 3 (3q) encompassing multiple cancer-associated genes, including *GMPS*,

*MLF1, TBL1XR1, PIK3CA, SOX2, and ETV5*. Additionally, an interstitial deletion on 20q, involving *ASXL1*, and a large terminal gain on 6q were detected. Notably, the analysis revealed a distinct region of copy-neutral loss of heterozygosity (CN-LOH) on 9p, which encompasses the critical cancer-associated genes *JAK2* and *CD274* (PD-L1).

#### Clinical Course

One week following the biopsy the patient was admitted to begin chemotherapy consisting of vincristine, dactinomycin, and cyclophosphamide [VDC] alternating with ifosfamide.

After completing one cycle of this treatment there was radiographic evidence of disease progression and oral crizotinib was added for indefinite maintenance. Six weeks from diagnosis palliative radiation of 45 Gy with an internal boost to 67.5 Gy to the primary tumor was undertaken after which he received two further cycles of VDC. Follow-up PET imaging revealed continued and rapid progression of pulmonary metastases. Chemotherapy was discontinued due to lack of response, and the patient passed away from disease 3.5 months after diagnosis.

### Establishment of a Patient-Derived Xenograft (PDX) Model and *In Vivo* Growth Kinetics

To establish a robust preclinical platform for investigating oncogenic signaling pathways and conducting functional drug sensitivity testing, a subcutaneous PDX model of the patient’s iRMS was successfully generated. Following the initial tissue implantation, the time to engraftment varied between the implanted tumor fragments. One xenograft (OHSU-BST-014D) exhibited a short latent phase of approximately 20 days with no palpable growth, followed by rapid exponential expansion that reached a volume of 2,000 mm^3^ by day 49. A second xenograft (OHSU-BST-014C) demonstrated a prolonged latent phase of approximately 100 days before entering a similar exponential growth phase (**Figure 4A**). This initial variability is consistent with the heterogeneous nature of primary tumor fragments. However, serial transplantation of the established PDX tumors into both nude (Nu/J) and NSG® immunocompromised mice subsequently demonstrated robust and consistent engraftment, indicating that aggressive tumor growth was maintained irrespective of the host’s specific extent of immune deficiency (Supplementary Figure 1).

**Figure 4.**
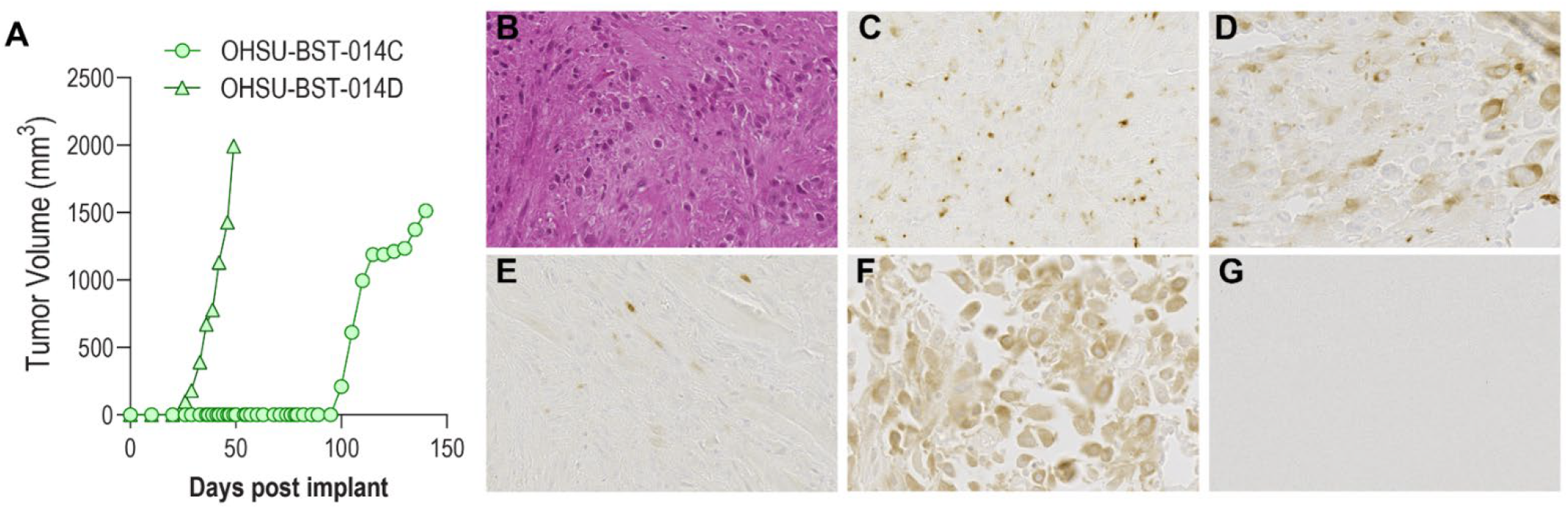
Establishment and characterization of the *FUS::TFCP2* iRMS patient-derived xenograft. (A) In vivo tumor growth kinetics of two engrafted fragments: OHSU-BST-014D and OHSU-BST-014C. (B) H&E staining of the PDX tumor and Immunohistochemistry for (C) desmin (diffuse cytoplasmic), (D) myogenin, (E) pan-keratin AE1/AE3, (F) ALK, (G) SATB2.

### Histologic and Immunohistochemical Analysis of the PDX Model

Hematoxylin and Eosin-stained section is morphologic features consistent with the original patient tumor (**Figure 4B)**. Immunohistochemistry showed diffuse cytoplasmic desmin expression (**Figure 4C**) and focal nuclear myogenin reactivity (**Figure 4D**). The tumor exhibited aberrant pankeratin (AE1/AE3) expression (**Figure 4E**) alongside strong ALK positivity (**Figure 4F**) and expected absence of SATB2 staining (**Figure 4G**). Together, these findings demonstrate that the established PDX tumors successfully preserved the primary diagnostic and immunohistochemical profiles observed in the original patient specimen.

### Molecular Characterization of the PDX Model

To validate that the established PDX preserved the molecular signature/characteristics of the primary malignancy, RT-PCR analysis was performed on cDNA synthesized from RNA isolated from PDX tumors. This analysis confirmed the *FUS::TFCP2* fusion transcript (Supplementary Figure 2A). We also evaluated the expression of *ALK* in the PDX using RT-PCR which confirmed expression of full length *ALK* transcript given that primers directed to extracellular and kinase domain both yielded expected amplicons (**Supplementary Figure 2B**). The pMIG ALK F1174L plasmid served as a positive control for the primers.

We generated a primary cell line from the PDX tumor (**Figure 5A**) and confirmed ALK protein expression by immunoblotting of cell lysates (**Figure 5B**). The blot revealed multiple ALK-immunoreactive species of differing molecular weight; lower-molecular-weight bands (~90 to 140 kDa) predominated, whereas the full-length protein at the predicted ~220 kDa was detected at lower intensity. The neuroblastoma cell line SH-SY5Y and the HEK293 cell line served as positive and negative ALK expression controls, respectively. Interestingly, while the primary (P1) culture was robust, we were unable to expand or passage this culture beyond this initial passage. Thus, all subsequent functional drug dose response experiments required seeding of PDX-tumor dissociated primary cells.

**Figure 5.**
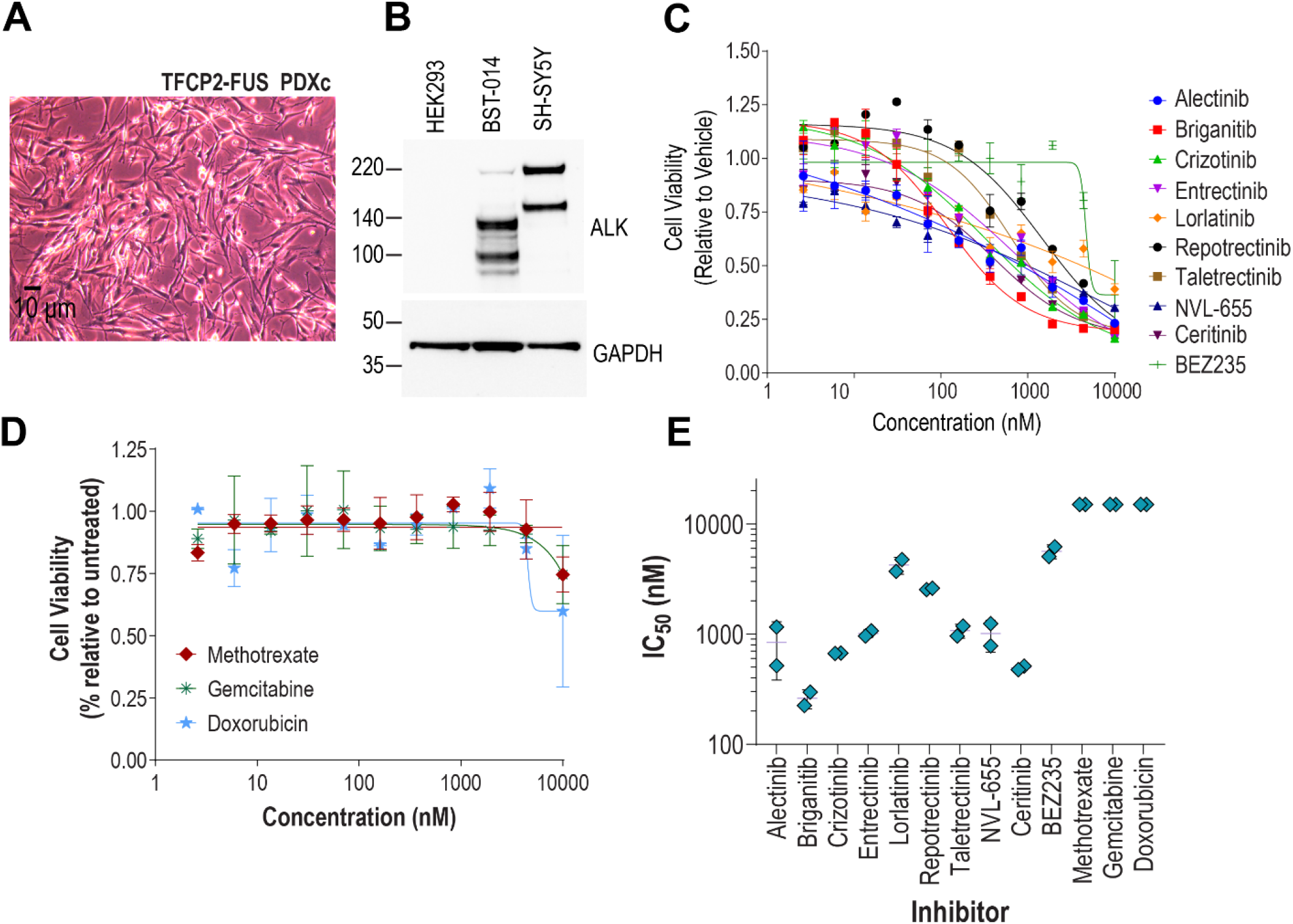
PDX-derived cell line and pharmacological sensitivity profiling. (A) Phase-contrast image of the PDX-derived primary cell line. (B) Immunoblot of PDX-derived cell lysates showing ALK expression. SH-SY5Y and HEK293 served as positive and negative controls, respectively. (C–D) Dose-response cell-viability curves across indicated agents. (E) Cell-based inhibitory concentration (IC_50_) for tested agents.

### Pharmacological Evaluation of ALK Inhibitors in PDX-Derived Cells

We performed an in vitro drug dose response assay on PDX-derived tumor cells and evaluated a panel of tyrosine kinase inhibitors (TKI) with demonstrable activity against ALK, including first-(crizotinib), second-(alectinib, brigatinib, and ceritinib), third-(lorlatinib), and fourth-generation (NVL-655) agents (Table S2), as well as the dual PI3K/mTOR inhibitor BEZ235 (dactolisib) for comparison. Chemotherapeutic agents (methotrexate, gemcitabine, and doxorubicin) were included to evaluate comparative efficacy against conventional cytotoxic regimens.

These experiments revealed varied sensitivity of the PDX-derived primary cells across the ALK TKI panel (**Figure 5C-E**). The sensitivity profile demonstrated a clear distinction between multi-kinase inhibitors and highly selective ALK inhibitors. Among the tested agents, the second-generation inhibitor brigatinib exhibited the most robust inhibitory activity with a cell-based 50% inhibitor concentration (IC_50_) of ~0.25 µM, followed closely by ceritinib (IC_50_ ~0.50 µM). The first-generation inhibitor crizotinib also demonstrated activity although the doses needed for efficacy approached 1 micromolar (IC_50_ ~0.70 µM).

In contrast, highly selective ALK inhibitors were less potent in these experiments. Alectinib and the highly potent and most selective fourth-generation inhibitor, NVL-655, exhibited only moderate activity (IC_50_ ~0.80 µM and ~1.0 µM, respectively), while lorlatinib failed to elicit a potent response (IC_50_ >4.0 µM). Other targeted agents, including the ROS1/pan-TRK inhibitors entrectinib, repotrectinib, and taletrectinib, showed IC50 values ranging from approximately 1.0 µM to 3.0 µM. The dual PI3K/mTOR inhibitor BEZ235 (dactolisib), with an IC_50_ of approximately 5.0 µM was also largely ineffective.

Conversely, the cytotoxic chemotherapeutic agents demonstrated negligible therapeutic efficacy, failing to reach an IC_50_ within the dose range where 10 µM was the highest concentration tested **(Figure 5D & 5E)**.

## Discussion

We report a 29-year-old man with metastatic pelvic iRMS harboring a *FUS::TFCP2* fusion and ALK overexpression. From this patient’s tumor, we established a PDX model that preserved the histologic, immunophenotypic, and molecular features of the primary tumor, providing a functional platform for direct, comparative pharmacological testing that genomics alone cannot supply. Two observations stand out: the tumor’s broad resistance to conventional chemotherapy, and an uneven response across ALK inhibitors that does not fit a conventional *ALK* fusion-driven model.

In *ALK* fusion-driven cancers, tumors are typically dependent on ALK kinase activity and respond to selective ALK inhibition^21,22^. The iRMS PDX, despite harboring ALK overexpression, behaved differently. The primary PDX-derived cells were sensitive to crizotinib, brigatinib, and ceritinib, yet resistant to the more ALK-selective potent TKI alectinib and lorlatinib, and to the highly selective and potent fourth-generation inhibitor NVL-655. This inhibitory profile where multi-kinase inhibitors with activity on ALK having higher cell-killing activity relative to more selective ALK TKI, points away from simple ALK dependence. Crizotinib inhibits MET, RON, and ROS1 in addition to ALK^23^; ceritinib additionally inhibits IGF1R and INSR^24^; and brigatinib has a broad profile that includes ROS1, FLT3, EGFR variants, IGF1R, and INSR^25^. The inactive drugs are progressively more ALK-selective, with NVL-655 shown to spare most of the kinome, including TRK^26^. Efficacy therefore tracked inversely with ALK selectivity, suggesting that activity in this tumor depends on co-inhibition of one or more kinases beyond ALK rather than on ALK blockade alone. Among the shared off-targets of the active agents, IGF1R and INSR are the most plausible single candidates; future studies are warranted.

These findings both align with and extend those of Schöpf and colleagues, who reported the ALK variants to be targetable, with ceritinib most effective, crizotinib intermediate, and the more ALK-selective alectinib least effective^14^. Clinically, however, ALK inhibitors have provided limited benefit in *TFCP2*-rearranged RMS; most reported patients have progressed, including rapid progression on crizotinib and on lorlatinib, with only rare and short-lived responses, such as a single near-complete response to lorlatinib lasting approximately six months^15^. This predominantly poor clinical experience is consistent with the resistance to selective ALK inhibitors that we observed in vitro, while the rare exceptions caution that in vitro sensitivity does not fully predict clinical response.

A second, independently derived preclinical model of TFCP2-rearranged intraosseous RMS, from a maxillary primary tumor, was recently reported by Bergsma and colleagues^27^. That model carried the CDKN2A/MTAP co-deletion and responded to combined PI3K/AKT and PRMT5 inhibition. Our PDX, from a pelvic primary, retained both loci and showed no such dependency; instead, we identified a distinct, ALK-centered polypharmacology. This raises the possibility that CDKN2A/MTAP status stratifies TFCP2-rearranged iRMS toward different actionable targets.

The ALK protein itself may be atypical here. Immunoblotting of PDX-derived cells revealed multiple ALK-immunoreactive species of differing molecular weight, with lower-molecular-weight bands predominating over full-length ALK. Schöpf et al. reported that *TFCP2*-rearranged sarcomas express several truncated ALK variants generated by intragenic deletions, alternative transcription, and aberrant splicing; these variants retain the kinase domain and are oncogenic and inhibitor-sensitive^14^. Our observed ALK banding pattern may be consistent with expression of such variants, although hypoglycosylated full-length ALK can also migrate below the mature receptor. Variant-specific differences in ALK conformation could plausibly contribute to the unusual inhibitor-sensitivity profile.

Transcript-level analysis, for example differential amplification across the ALK extracellular and kinase domains, which were not performed in this study, could confirm the presence of truncated variants directly.

The tumor’s resistance to methotrexate, gemcitabine, and doxorubicin is consistent with the poor chemo-responsiveness reported for *TFCP2*-fused RMS^15^. Because single-agent selective ALK inhibition was ineffective, strategies that engage additional dependencies, whether through multi-target inhibitors or rational combinations, warrant systematic evaluation in this entity.

Cytogenetic profiling identified copy-neutral LOH of 9p, a region that includes *JAK2* and *CD274* (PD-L1). Copy-neutral LOH can reinforce an existing driver by duplicating a mutant allele while removing its wild-type counterpart^28,29^. Here, however, no activating *JAK2* mutation was detected, and the event is copy-number neutral, so it provides no dosage gain at *CD274*; the functional significance of this LOH is therefore unclear. Notably, and in contrast to the recurrent *CDKN2A/MTAP* co-deletion reported in this entity^14^, both loci were intact in our case, and a PRMT5-directed rationale therefore did not apply.

Several limitations apply. Our conclusions derive from a single patient and a single PDX. The in vitro pharmacology was not confirmed by in vivo efficacy testing, and the attribution of activity to specific kinases is inferential, drawn largely from published selectivity profiles obtained in other tumor contexts. The primary cells could not be expanded in vitro, beyond early passage, which limited independent re-testing in these cell lines; the same constraint also highlights that the drug screen was performed on minimally cultured cells that closely reflect the tumor of origin.

In conclusion, here we demonstrate that *FUS::TFCP2* iRMS tumors cells are more responsive to multi-target rather than ALK-selective TKI, a pattern that should temper expectations for narrow-spectrum ALK-directed therapy in these cases. Defining the responsible co-target(s) and confirming activity in vivo are the key next steps, and the model reported here provides a tractable platform for both.

## Acknowledgements

This project was supported by funds from the Kuni Foundation (Portland, Oregon).

