## Supplementary Figures and Table for "A patient-derived xenograft model of *FUS::TFCP2* intraosseous rhabdomyosarcoma reveals chemoresistance and differential sensitivity to ALK inhibitors"

### Supplementary Materials

### Supplementary Figure 1

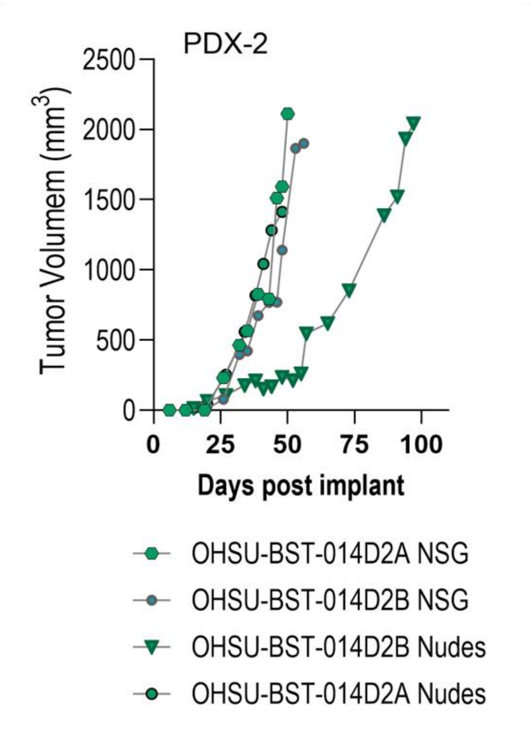


Serial transplantation of OHSU-BST-014 in nude (Nu/J, Jackson Lab Strain Strain #:002019) and NSG® Jackson Lab Strain Strain #:005557) immunocompromised mice.

**Supplementary Figure 1.** Serial transplantation of OHSU-BST-014 in nude (Nu/J, Jackson #002019) and NSG (Jackson #005557) immunocompromised mice.

**Supplementary Figure 2**

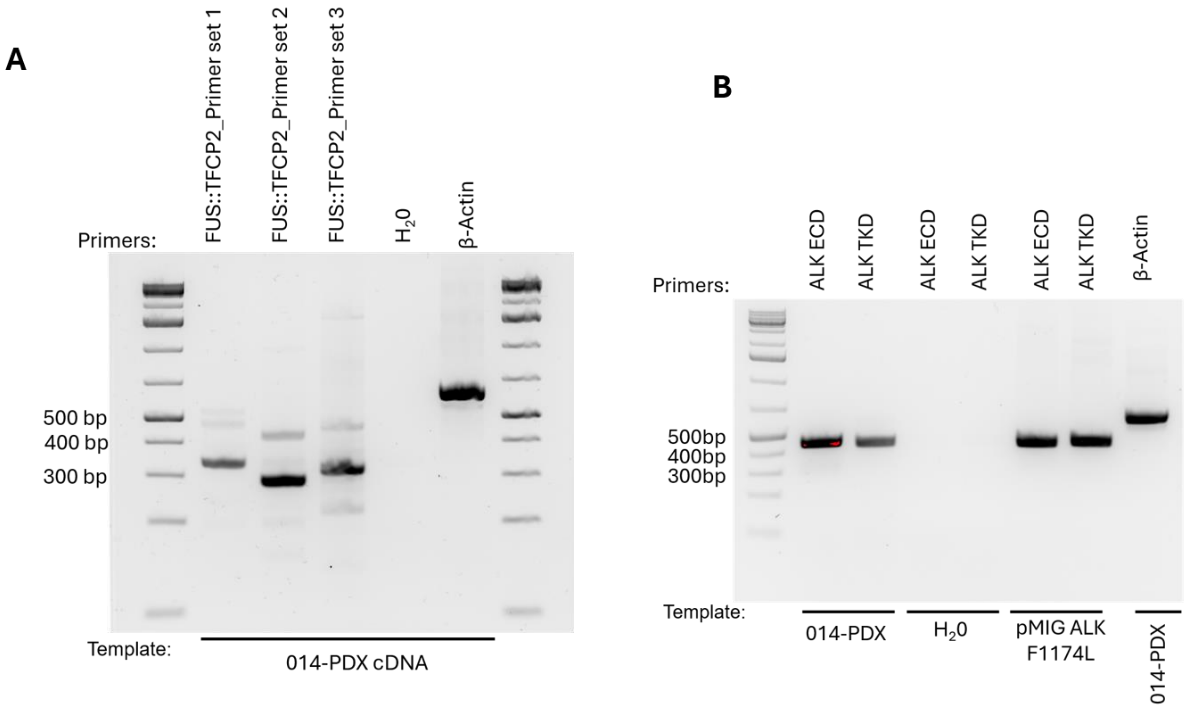


**Supplementary Figure 2.** (A) RT-PCR confirming the *FUS::TFCP2* fusion transcript in PDX tumor cDNA. (B) RT-PCR confirming full-length *ALK* transcript, with amplicons from both extracellular-domain and kinase-domain primers; the pMIG ALK-F1174L plasmid served as a primer positive control.

**Table S1.** The sequence of the set of primers used for RT-PCR to detect the fusion protein, FUS::TFCP2.

| **Primer Name** | **Primer Sequences** | | **Amplicon** |
|---|---|---|---|
| | **Forward** | **Reverse** | |
| **FUS::TFCP2-1** | CCAAGATCAATCCTCCATGAGTAG | CTTTGTGCTGCTACCTCTCC | 333 bp |
| **FUS::TFCP2-2** | GCGGTTATGGCAATCAAGAC | AATATGTGCTTTGTGCTGCTAC | 286 bp |
| **FUS::TFCP2-3** | GCAGTGGTGGCGGTTAT | CTACCTCTCCAGCAGTGAAAC | 312 bp |
| **ALKECD1** | GAAGAGTCTGGCAGTTGACTTC | GAAGTGGCCAGAGAGGCAAG | 456 bp |
| **ALKTKD1** | CTGCAAGTGGCTGTGAAGAC | GAATTGACCGTCGTACCGTC | 450 bp |
